# Gut feeling: Extent of virulence and antibiotic resistance genes in *Helicobacter pylori* and campylobacteria

**DOI:** 10.1101/2023.09.27.559685

**Authors:** R. Shyama Prasad Rao, Sudeep D. Ghate, Larina Pinto, Prashanth Suravajhala, Prakash Patil, Praveenkumar Shetty, Nagib Ahsan

## Abstract

**Background:** *Helicobacter pylori*, a member of campylobacteria, is the leading cause of chronic gastritis and gastric cancer. Virulence and antibiotic resistance of *H. pylori* are of great concern to public health. However, the relationship between virulence and antibiotic resistance genes in *H. pylori* in relation to other campylobacteria remains unclear.

**Materials and Methods:** By using the virulence and comprehensive antibiotic resistance databases, we explored all available 354 complete genomes of *H. pylori* and compared it with 90 species of campylobacteria for virulence and antibiotic resistance genes/proteins.

**Results:** On average, *H. pylori* had 129 virulence genes, highest among *Helicobacter spp*. and 71 antibiotic resistance genes, one of the lowest among campylobacteria. Just 2.6% of virulence genes were shared by all campylobacterial members, whereas 9.4% were unique to *H. pylori*.

The cytotoxin-associated genes (cags) seemed to be exclusive to *H. pylori*. Majority of the isolates from Asia and South America were *cag2*-negative and many antibiotic resistance genes showed isolate-specific patterns of occurrence. Just 15 (8.8%) antibiotic resistance genes, but 103 (66%) virulence genes including 25 cags were proteomically identified in *H. pylori*.

Arcobacterial members showed large variation in the number of antibiotic resistance genes and there was a positive relation with the genome size.

**Conclusion:** Large repository of antibiotic resistance genes in campylobacteria and a unique set of virulence genes might have important implications in shaping the course of virulence and antibiotic resistance in *H. pylori*.

## Introduction

*Helicobacter pylori* is a gram-negative microaerophilic bacterium that persistently colonizes the epithelial lining of the stomach, and it is estimated that more than half the world population is chronically infected (Hooi et al., 2017). *H. pylori* has been associated with numerous pathophysiological conditions including chronic gastritis, peptic ulcer, gastric cancer, and gastric mucosa-associated lymphoid tissue lymphoma (Suerbaum and Michetti, 2002). Annual stomach cancer rates are high – there were more than 1.22 million incident cases and 865000 deaths in 2017 that contributed to 19.1 million disability-adjusted life-years (Etemadi et al., 2020). *H. pylori* infection is the most important established risk factor for stomach cancer (Chiang et al., 2021; Choi et al., 2020; Malfertheiner et al., 2017). The more virulent *cagA*-positive *H. pylori* strains also lead to persistent inflammatory stimulation and have been associated with ischemic heart disease (Pasceri et al., 1998).

Given the high burden of *H. pylori*, especially in the developing countries (Etemadi et al., 2020; Hooi et al., 2017), elimination of *H. pylori* infection has been suggested and attempted (Chiang et al., 2021; Choi et al., 2020). *H. pylori* eradication quadruple therapy usually includes the heavy metal bismuth, a proton pump inhibitor such as lansoprazole, and at least two antibiotics among amoxicillin, clarithromycin, furazolidone, levofloxacin, metronidazole, and tetracycline (Hu et al., 2017). However, as *H. pylori* resistance to many of these antibiotics is already at alarming levels worldwide (Savoldi et al., 2018), the increasing prevalence of antibiotic resistance in *H. pylori* has been recognized by the WHO as “high priority” for urgent need of new therapies (Tacconelli et al., 2018).

There are attempts at finding novel and effective therapeutic regimens for *H. pylori* (Hu et al., 2017), including antibiotics susceptibility-guided treatment (Gingold-Belfer et al., 2021). However, as *H. pylori* seems to be resistant to multiple antibiotics such as clarithromycin, levofloxacin, metronidazole, and tetracycline at high concentration (Megraud et al., 2021; Nestegard et al., 2022; Shu et al., 2022), there is a need to understand the antibiotic resistance in *H. pylori* from different perspectives. Two areas standout – first one is the relationship between the virulence and antibiotic resistance genes within *H. pylori*, and the second one is the extent of virulence and antibiotic resistance genes between *H. pylori* and other campylobacteria.

Recent studies are revealing some interesting relationships between virulence and antibiotic resistance. For instance, while Brennan et al. (2018) found that less virulent (*cagA*-negative and *vacA* S2-containing) strains of *H. pylori* are associated with primary clarithromycin resistance, da Silva Benigno et al. (2022) concluded that virulent *H. pylori* strains (*cagA* and *cagE* positive) may be more susceptible to clarithromycin treatment. However, Hosseini et al. (2021) found that isolates with virulence genes *oipA, vacA*, and *iceA1* were resistant to clarithromycin, while Liu et al. (2022) found no relationship between the presence of virulence factors *cagA* and *vacA*, and resistance to clarithromycin, levofloxacin, and metronidazole.

The comparison of *H. pylori* with other campylobacteria is also very relevant and useful for multiple reasons. Many campylobacterial members are zoonotic and similar to *H. pylori* they cause gastroenteritis. Further, apart from being also associated with human extragastrointestinal infections, *Campylobacter spp*. are also the leading cause of bacterial foodborne and waterborne infections (Igwaran and Okoh, 2019). In fact, *Campylobacter* infection is widespread, and the incidence and prevalence of campylobacteriosis have been increasing (Kaakoush et al., 2015). For example, *Campylobacter concisus* gastritis was often misattributed to *H. pylori* on gastric biopsy (Ferreira et al., 2022). *Campylobacter spp*. infection and antibiotic resistance are widespread especially in sub-Saharan Africa (Hlashwayo et al., 2021). The fluoroquinolone-resistant *Campylobacter spp*. are recognized by the WHO as “high priority” (Tacconelli et al., 2018), and campylobacteriosis was the most reported zoonosis in the European Union in 2020 (EFSA, 2022).

While the knowledge on the relationship between virulence and antibiotic resistance genes within *H. pylori* is sparse and uncertain (Brennan et al., 2018; da Silva Benigno et al., 2022; Hosseini et al., 2021; Liu et al., 2022; Wang et al., 2019), the extent of virulence and antibiotic resistance genes between *H. pylori* and other campylobacteria is largely unexplored.

In this study, we sought to investigate the relationship between virulence and antibiotic resistance genes among all available 354 complete genomes of *H. pylori* and also wanted to get a larger perspective by comparing it with 90 species of other campylobacteria. We used the comprehensive antibiotic resistance database (CARD) (Alcock et al., 2019) and virulence factor database (VFDB) (Liu et al., 2019) – two most inclusive and widely used resources for the genome-wide identification of virulence and antibiotic resistance genes, respectively. We reveal interesting patterns of virulence and antibiotic resistance genes among campylobacteria and *H. pylori*, and discuss their relevance in shaping the scope of antibiotic resistance.

## Materials and Methods

### Genome sequence acquisition

The complete genome reference sequences (RefSeqs) for all campylobacterial species were downloaded from the NCBI website (https://www.ncbi.nlm.nih.gov/, last accessed on Dec 15, 2022). There were 91 species (from 15 genera) of campylobacteria with a RefSeq. Further, all available 353 complete genome sequences (excluding RefSeq) for *H. pylori* were also downloaded. For most of the other campylobacterial species, apart from RefSeq, the NCBI database did not contain additional complete genome sequences. Strict filter criteria were used to ensure the quality as only annotated genomes with an assembly level of “complete” were used. They were also confirmed to have a single circular chromosome with the size matching the average genome size of the species. Genome sequences were downloaded in “.fna” file format. Based on the coordinate information available in the corresponding “.gtf” file, corresponding 16S rRNA sequences from the above genomes were used for phylogenetic analysis. All sequence related metadata including the geographical origins of the isolates were also collected from the NCBI. All relevant basic information about the genomes such as length, GC content (%), geographical origins of the isolates, etc. are given in Table S1.

### Identification of virulence and antibiotic resistance genes

Numerous databases such as AMRFinderPlus (Feldgarden et al., 2021), ARDB (Liu and Pop, 2009), CARD (Alcock et al., 2019), MEGARes (Doster et al., 2020), and ResFinder (Bortolaia et al., 2020), exist for the computational identification of resistance determinants in the genomes (Hendriksen et al., 2019). However, each one has limitations. For example, ARDB (https://ardb.cbcb.umd.edu/) is not maintained anymore and MEGARes (https://www.meglab.org/megares/) redirects to CARD. A comparison of AMRFinderPlus (developed by NCBI, https://www.ncbi.nlm.nih.gov/pathogens/antimicrobial-resistance/AMRFinder/) prediction with CARD prediction showed that the former result was only a subset of the later (Rao et al., 2023). Thus, the comprehensive antibiotic resistance database (CARD, https://card.mcmaster.ca/) is the most inclusive tool for the prediction of antibiotic resistance genes (ARG) (Liu et al., 2022). As defined in CARD, ARGs are the molecular determinants – genes and their regulators conferring resistance to antibiotic molecules. The gene sequences and their mutations, if any, are known, and clear experimental evidence exists (Alcock et al., 2019). As a curated data resource, the CARD has 4336 antibiotic resistance ontology (ARO) terms for 2923 known antimicrobial resistance (AMR) determinants/genes and 1304 resistance variant mutations (Alcock et al., 2019). The web interface of CARD (https://card.mcmaster.ca/analyze/rgi) identifies putative antibiotic resistance genes from experimentally confirmed AMR gene models based on multiple approaches such as BLAST, sequence alignment, regular expressions (RegEx), hidden Markov models (HMMs), and/or position-specific SNPs (Alcock et al., 2019). We used the web interface with RGI version 6.0.1 and CARD 3.2.7. Further, we used the complete genomic DNA sequence of high quality/coverage and the prediction of partial genes were excluded. Each campylobacterial genome sequence was submitted to CARD’s resistance gene identifier (RGI) tool to obtain annotations based on perfect, strict, or loose paradigm, and complete gene match criteria for the identification of antibiotic resistance genes (Rao et al., 2023; Zhang et al., 2022). The CARD outputs a list of ARO terms – the total number of antibiotic resistance genes – for each genome. Duplicate (two or more) entries, if any, of the ARO term indicate multiple copies of an antibiotic resistance gene (Rao et al., 2023).

The virulence factor database (VFDB, http://www.mgc.ac.cn/VFs/) is a comprehensive warehouse and online platform widely used for the identification of virulence factors (VFs) (Liu et al., 2019; 2022). The integrated and automatic pipeline VFanalyzer (http://www.mgc.ac.cn/cgi-bin/VFs/v5/main.cgi) in the VFDB systematically identifies known/potential VFs in complete genomes based on iterative and exhaustive sequence similarity searches of orthologous groups within the query genome and preanalyzed reference genomes from VFDB to eliminate potential false positives from paralogs (Liu et al., 2019). We used Abricate 0.9.8 (https://github.com/tseemann/abricate) interface to screen the VFDB using the parameters – percentage identity of ≥60□%□and coverage of ≥40□% (Mootapally et al., 2021).

### Phylogenetic analysis

To understand virulence and antibiotic resistance genes among campylobacteria from a phylogenetic perspective, complete 16S rRNA sequences from 91 species of campylobacteria were used to construct a phylogenetic tree in MEGA11 using the maximum likelihood method (Tamura et al., 2021). A thousand bootstrap iterations were performed. *Nautilia profundicola* was taken as the outgroup.

### Data analyses

The total and unique number of genes were counted for individual species and the extent of overlap of genes among different species/clades were represented using a Venn diagram (Rao et al., 2023). The R function/package ggvenn() was used to make the Venn diagram. Scatter plot was used to visualize the relationship between two sets of data, for example, the number of virulence genes versus the number of antibiotic resistance genes. To account for the different sizes of genomes, the number of genes were normalized by dividing with individual genome size (in bp) and multiplied by the average genome size. The Spearman’s rank correlation, which is less sensitive to outliers, was used to examine the relationship between the numbers of antibiotic resistance genes and virulence genes. The significance of the correlation coefficient was tested using cor.test (which is based on t-distribution or approximation) in R. As the extent of antibiotic resistance is also associated with the differential presence of virulence genes, in particular cytotoxin-associated genes (cags) (Brennan et al., 2018; da Silva Benigno et al., 2022; Hosseini et al., 2021; Liu et al., 2022), to further understand their interplay, genomes were grouped based on cags into three categories namely “all cags” when all cags were present, “*cag2-*” when only *cag2* was missing, and “others” when most of the cags including *cag2* and *cagA* were missing.

An unpaired t-test (two-tailed, unequal variance) was used to compare the means (for instance, the average number of genes) between two groups (Rao et al., 2023). In addition, a chi-squared test was used (as chisq.test() in R with the parameter simulate.p.value = TRUE) for the analysis of contingency table, for example, to check whether the difference in the numbers of genes/genomes in different categories was significantly different (Agresti, 2018). A Bonferroni correction was applied to account for multiple comparisons. The extent of overlap of genes between two sets was quantified using overlap coefficient which is defined as the size of the intersection of two sets divided by the size of the smaller of the two sets (Vijaymeena and Kavitha, 2016). Finally, to see if virulence and antibiotic resistance genes were proteomically identified, we searched the literature for proteomic studies using PubMed and Google Scholar with keywords such as “*Helicobacter pylori*” AND “proteomics” (Karlsson et al., 2016). The overlaps between different sets were depicted using a Venn diagram. The routine data handling/analysis was done in Python and Microsoft Excel.

## Results

### Number of antibiotic resistance and virulence genes in H. pylori and campylobacteria

The *H. pylori* and other campylobacterial complete genomes were explored using CARD and VFDB for the identification of antibiotic resistance and virulence genes, respectively (Table S2-S9). There were, on average, 90.9 (SD ±2.0, range 86-95) antibiotic resistance genes and 132.8 (SD ±11.0, range 104-155) virulence genes in the genomes of *H. pylori* (Table 1). As a few genes were in multiple copies, after ignoring those duplicate entries, there were 71.4 (SD ±2.2, range 66-77) unique antibiotic resistance genes and 128.9 (SD ±10.9, range 101-145) unique virulence genes. It should be noted that CARD annotates antibiotic resistance genes based on protein homolog and variant models, and perfect/strict/loose paradigm. For “loose” hits, the median bit score of 99 (average = 158.5) with median sequence length of 1074 (average = 1309.7) would translate to an E-value of 7.35E-24 (1.1E-41 for average). In case of virulence genes, majority of the hits had far higher sequence identity and coverage than the assigned cut-offs of ≥60□%□and ≥40□%, respectively. For example, 92.8% hits had ≥85% of sequence identity and ≥70% of coverage. The lowest coverage sequence at 40.04%, with a length of 473 and 69.86% of identity, had an E-value of 2.44E-51. Whereas the lowest identity sequence at 63.47%, with a length of 1095 and 54.11% of coverage, had an E value of 1.02E-121. Compared to *H. pylori*, other species of *Helicobacter* had slightly higher number of antibiotic resistance genes (82.5±9.3, 68-98, p = 0.001, t-test), but only half the number of virulence genes (65.5±20.8, 34-109, p = 1.1E-07, t-test). The *Wolinella*, a close relative of *Helicobacter* (Fig. S1), had just 28 virulence genes, but 115 antibiotic resistance genes with some of them in multiple copies (Table S6). Compared to *Helicobacter spp*., *Campylobacter spp*., on average, had similar number of antibiotic resistance genes (88.2±6.7, 77-106, p = 0.057, t-test), but more virulence genes (86.3±40.4, 9-155, p = 0.029, t-test). Arcobacterial species and *Sulfurospirillum spp*., on average, had far lower number of virulence genes, but had much higher number of unique antibiotic resistance genes with many in numerous copies that nearly doubled their total number (Table 1 and S6).

**Table 1.** Summary of virulence and antibiotic resistance genes in campylobacteria.

| Clade | # of species | # of antibiotic resistance genes<sup>&</sup> |  | # of virulence genes<sup>&</sup> |  |
| --- | --- | --- | --- | --- | --- |
|  |  | Unique | Total | Unique | Total |
| *H. pylori*<sup>+</sup> | 1 | 71.4 (66-77) | 90.9 (86-95) | 128.9 (101-145) | 132.8 (104-155) |
| *Helicobacter spp.* | 13 | 82.5 (68-98) | 106.7 (84-136) | 65.5 (34-104) | 66.8 (34-109) |
| *Wolinella* | 1 | 115 | 204 | 28 | 28 |
| *Sulfuricurvum* | 1 | 134 | 254 | 18 | 21 |
| *Sulfurimonas spp.* | 9 | 107.9 (103-113) | 180.9 (144-206) | 29.4 (14-41) | 29.6 (14-41) |
| Arcobacterial species | 25 | 126.4 (94-156) | 255.6 (134-398) | 24.9 (18-33) | 26.0 (18-34) |
| *Campylobacter spp.* | 32 | 88.2 (77-106) | 124.1 (107-155) | 86.3 (9-155) | 88.8 (9-157) |
| *Sulfurospirillum spp.* | 5 | 134.0 (119-151) | 262.0 (215-309) | 37.8 (32-41) | 38.6 (32-42) |
| Others | 4 | 104.3 (91-114) | 162.5 (138-178) | 13.8 (6-22) | 14.3 (7-23) |
| All | 91 | 103.6 (68-156) | 174.6 (84-398) | 54.1 (6-155) | 55.6(7-157) |

<sup>&</sup>Average (min-max), <sup>+</sup>Based on 354 complete genomes for *H. pylori*. See Fig. S1 for a phylogenetic relationship between the clades.

### Comparison of antibiotic resistance and virulence genes among campylobacteria

Together, there were 613 antibiotic resistance genes and 446 virulence genes among 91 species (based on RefSeqs) of campylobacteria (Table S6 and S8). Whereas in 354 complete genomes of *H. pylori*, altogether, there were 170 antibiotic resistance genes and 156 virulence genes (Table S7 and S9). Based on Venn, 80 (∼12.8%) antibiotic resistance genes were common to all clades whereas only 16 (∼2.6%) were unique to *H. pylori* (Fig. 1A). The variants of *rpoB* and *gyrB*, for instance, were specific to *H. pylori*. In contrast, just 12 (2.6%) virulence genes were common to all clades whereas 43 (9.4%, primarily cags) were unique to *H. pylori* (Fig. 1B). There were substantial overlaps between Helicobacteraceae (including *H. pylori*) and Campylobacteraceae – 160 antibiotic resistance genes (overlap coefficient = 0.39) and 139 virulence genes (overlap coefficient = 0.50) were common to both. But there were also numerous unique genes in each family. The “others” set – a clade containing *Hydrogenimonas, Nitratifractor*, and *Sulfurovum* (Fig. S1) had nine (∼4.6%) unique antibiotic resistance genes and 4 (∼10.8%) unique virulence genes. Within Helicobacteraceae, *Wolinella* had very few unique antibiotic resistance and virulence genes (Fig. S2).

**Fig. 1.**
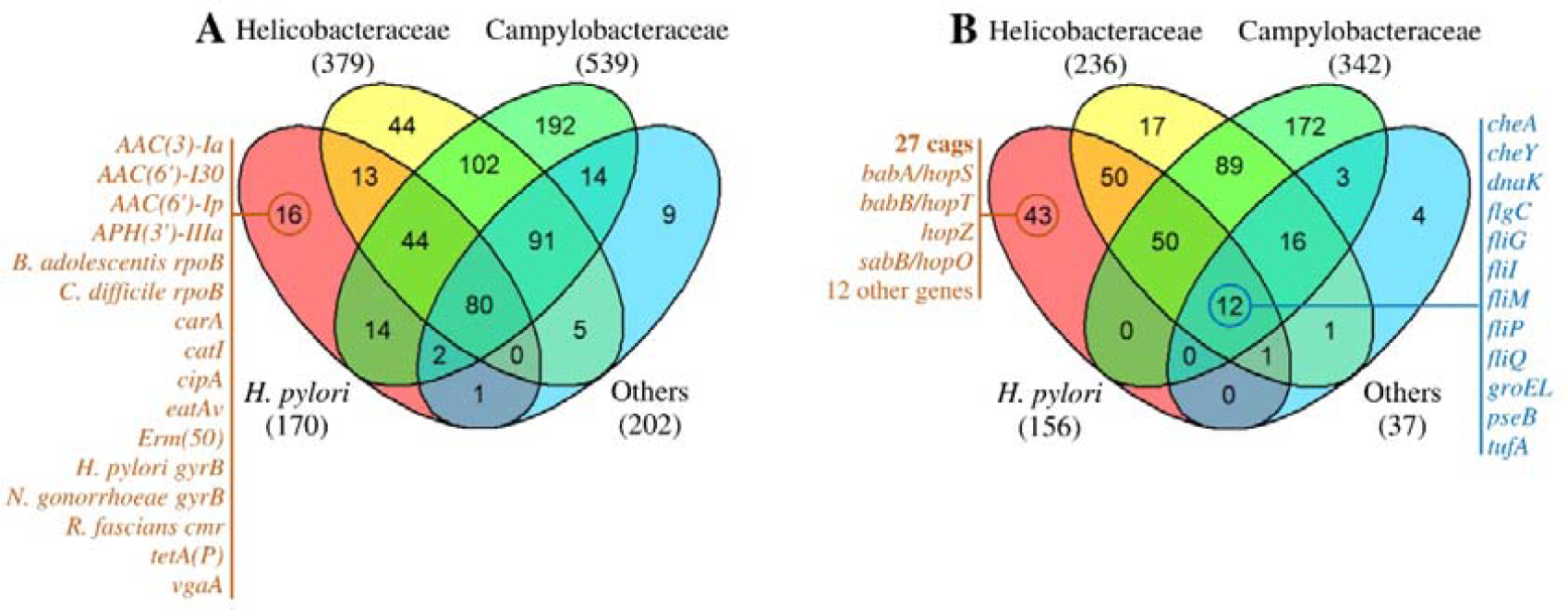
Venn diagrams showing the overlap in the presence of (A) antibiotic resistance genes and (B) virulence genes in campylobacterial clades. (A) Eighty (12.8%) antibiotic resistance genes were common to all clades while only 16 (2.6%) were unique to *H. pylori*. (B) Just 12 (2.6%) virulence genes were common to all clades whereas 43 (9.4%, primarily cags) were unique to *H. pylori*. Note: *H. pylori* was not included in the Helicobacteraceae set.

### Relationship between antibiotic resistance and virulence genes

There was a negative correlation (ρ = -0.64, p = 1.3E-11, t-test for correlation) between the total number of antibiotic resistance genes and the total number of virulence genes among the 91 species of campylobacteria (Fig. 2A). Further, species from different clades/genera formed distinct clusters based on the numbers of antibiotic resistance and virulence genes. Individually, the numbers of virulence genes were negatively correlated (ρ = -0.68, p = 1.3E-13, t-test for correlation) with the genome size, whereas the numbers of antibiotic resistance genes were strongly positively correlated (ρ = 0.91, p = 2.2E-16, t-test for correlation, Fig. 2B and 2C). Except for *Campylobacter* clade, the overall patterns remained similar after adjusting to the genome size.

**Fig. 2.**
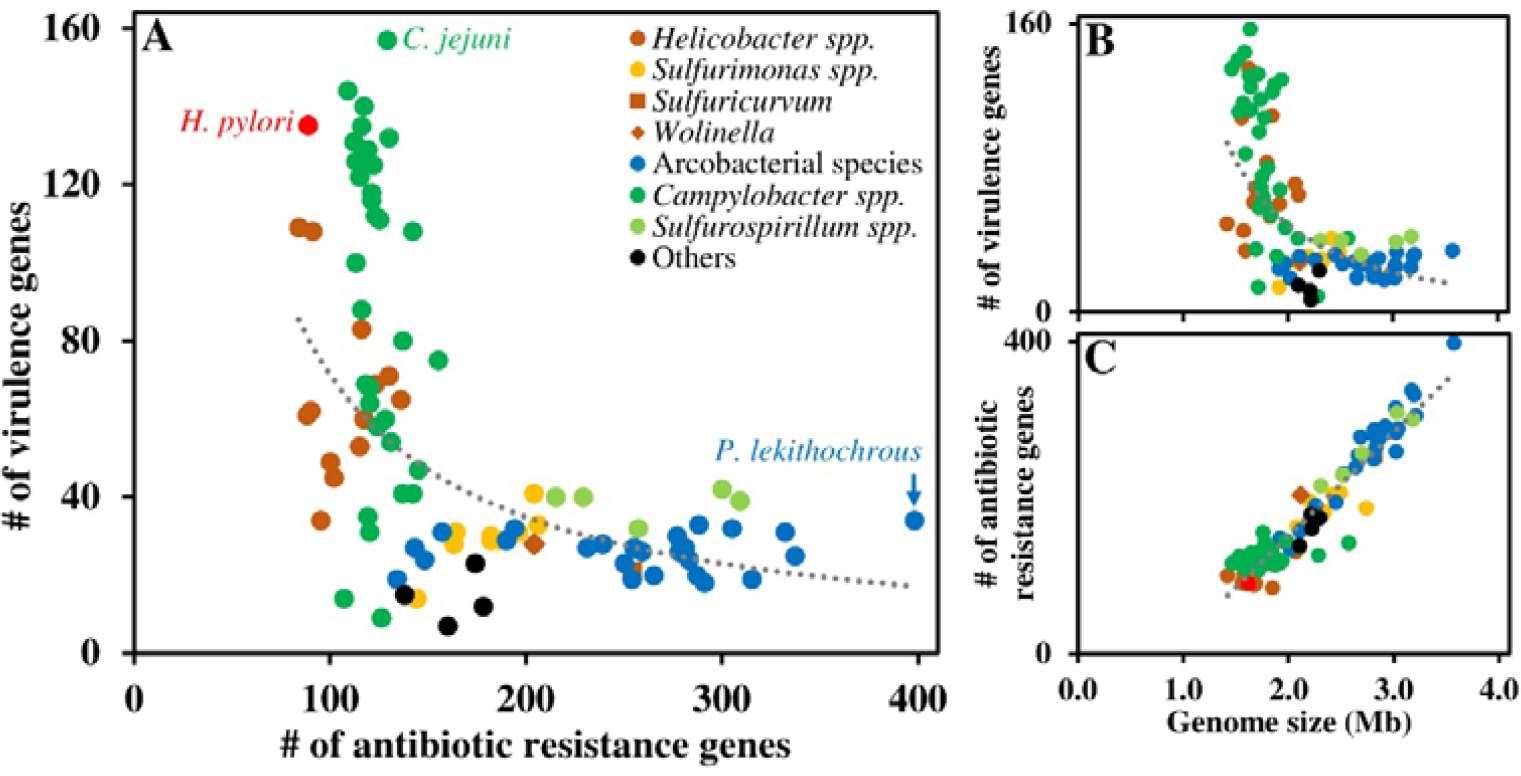
(A) Overall, the total numbers of antibiotic resistance genes were negatively correlated (ρ = -0.64) with the total numbers of virulence genes in campylobacteria. Species from different clades/genera form distinct clusters based on the number of virulence and antibiotic resistance genes. Further, (B) the number of virulence genes were negatively correlated (ρ = -0.68) with the genome size, whereas (C) the antibiotic resistance genes were positively correlated (ρ = 0.91).

### Antibiotic resistance versus virulence genes in H. pylori

There was a weak negative correlation (ρ = -0.15, p = 3.96E-3, t-test for correlation) between the numbers of antibiotic resistance genes and virulence genes in *H. pylori* (Fig 3A). However, genomes could be grouped based on the presence of cags – in 133 (37.6%) genomes all cags were present, in 147 (41.5%) genomes only *cag2* was absent, whereas in the remaining 74 (20.9%) genomes many or all cags were missing. Interestingly, the *cag2*-negative *H. pylori* genomes were almost all exclusive to Asia and South America (Fig. 3B). Further, apart from all cags, a few more virulence genes such as *htpB, groEL*, etc. were also significantly different (p < 0.05, chi-squared test with Bonferroni correction) in one of the groups based on cags (Fig. S3). Similarly, there were at least 25 antibiotic resistance genes that were preferentially present among these three groups (Fig. 3C, Table S7 and S9). Many of those genes were known to impart antibiotic resistance as *H. pylori* is resistant to numerous antibiotics (Table S10) (Alcock et al., 2019; Megraud et al., 2021; Nestegard et al., 2022; Savoldi et al., 2018; Shu et al., 2022). For instance, different variants of *rpoB* which impart resistance to rifampicin were present in different groups based on cags (Fig. 3C and Table S10).

**Fig. 3.**
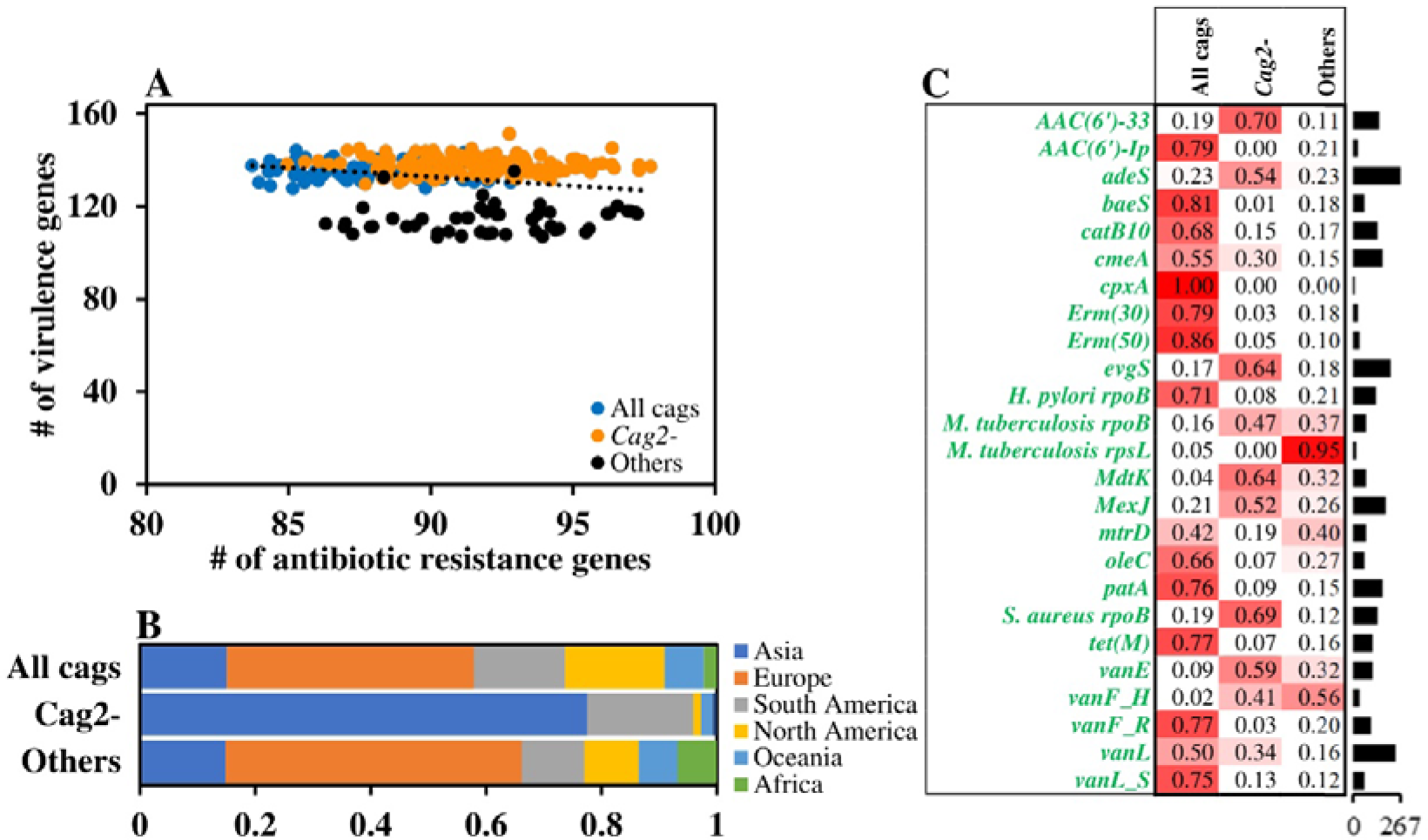
Antibiotic resistance versus virulence genes in *H. pylori*. (A) Correlation between the numbers of antibiotic resistance vs virulence genes was low (ρ = -0.15). (B) The *cag2*-negative *H. pylori* genomes were exclusive to Asia and South America. (C) Heat map shows the relative proportion of genes in three groups – at least 25 antibiotic resistance genes were preferentially present (p < 0.05, chi-squared test with Bonferroni correction) in one of the groups based on cags. Bar graph at the right shows the number of genomes.

Finally, we looked at the proteomic studies (Table S11) (Karlsson et al., 2016; Kumar et al., 2022; Loh et al., 2021; Müller et al., 2013; 2015; Sugiyama et al., 2019; Wei et al., 2022) to check if antibiotic resistance and virulence genes were proteomically known. Just 15 (8.8%) antibiotic resistance genes (such as *gyrA, gyrB*, and *rpoB*), but 103 (66.0%) virulence genes including all cags expect *cag2* and *cagP* were proteomically identified based on *H. pylori* proteomics (Fig. S4). This is because the *H. pylori* genomes were more diverse with respect to antibiotic resistance genes – only 67 (39.4%) antibiotic resistance genes but 128 (82.1%) virulence genes were present in 50% or more of genomes.

## Discussion

Given the virulent nature of *H. pylori* (Censini et al., 1996), it has a high global prevalence and disease burden (Hooi et al., 2017). In addition, the increasing antibiotic resistance in *H. pylori* is slowing its eradication rate (Hu et al., 2017; Megraud et al., 2021; Nestegard et al., 2022; Shu et al., 2022; Tacconelli et al., 2018). Further, given that numerous other members of campylobacteria are also pathogenic and antibiotic resistant (EFSA, 2022; Ferreira et al., 2022; Hlashwayo et al., 2021; Igwaran and Okoh, 2019; Kaakoush et al., 2015; Tacconelli et al., 2018), it is highly appropriate to look at the virulence and antibiotic resistance in *H. pylori* in the backdrop of entire campylobacterial clade.

*H. pylori* and *C. jejuni* in particular seem to have the highest number of virulence genes among 91 species of campylobacteria. Further, along with the numerous unique virulence genes, cags seemed to be exclusive to *H. pylori*. This hints at the high virulence and global prevalence of *H. pylori* and *C. jejuni* (EFSA, 2022; Hooi et al., 2017; Kaakoush et al., 2015; Tacconelli et al., 2018). The others, more specifically the arcobacterial members, showed low numbers of virulence genes. Most member of this clade are free living in a wide range of habitats, although a few are considered as emergent enteropathogens and/or potential zoonotic agents (Collado et al., 2011; Pérez-Cataluña et al., 2018). On the contrary, while *H. pylori* had one of the lowest numbers of antibiotic resistance genes, arcobacterial members showed a wide range with many containing several hundreds. It may be noted that the members of arcobacteraceae are very diverse and their hosts/habitats include aquatic animals and planktons, cyanobacterial mats, sludge and marine sediments, estuarine and river water, etc. (Pérez-Cataluña et al., 2018). It is believed that free-living bacteria acquire resistance factors to counter environmental challenges including antibiotic pollution and biotic hostility such as cyanobacterial bloom (Larsson et al., 2022; Zhang et al., 2020).

It is well known that *H. pylori* contains as many as 32 cags that encode a bacterial type IV secretion system, and it was observed that cags were present in approximately 60-70% of Western *H. pylori* strains and virtually 100% of East-Asian *H. pylori* strains (Noto et al., 2012). The CagA protein as a pro-oncogen requires tyrosine phosphorylation at EPIYA motif and structural polymorphism of CagA influences its scaffold function, which was thought to underlie the geographic difference in the incidence of gastric cancer (Hatakeyama et al., 2017). We found that the majority of the isolates from Asia and South America were *cag2*-negative. The *hp0521* (*cag2*) gene encodes Cag2 protein. Xu et al. (2022) have found that the deletion of *cag2* gene had no effect on bacterial growth but increased *cagA* mRNA expression and CagA protein thereby likely affecting the virulence of *H. pylori*. This may partly explain why most Asian strains of *H. pylori* are more virulent (Hatakeyama et al., 2017;Yuan et al., 2017).

The interplay between virulence and antibiotic resistance is documented – for example, *Pseudomonas aeruginosa* with *oprD* mutants that impart carbapenem resistance were also more virulent (Geisinger and Isberg, 2017). The *H. pylori* strains with virulence genes *oipA, vacA*, and *iceA1* were resistant to clarithromycin (Hosseini et al., 2021). On the contrary, there might also be a trade-off between the two – less virulent (*cagA*-negative) strains of *H. pylori* were found to be clarithromycin resistance (Brennan et al., 2018) and vice versa (da Silva Benigno et al., 2022). The differential presence of antibiotic resistance genes among *cag2*-positive and negative strains observed in this study further hints at such an interplay and trade-off.

Further, as *H. pylori* is known to be resistant to numerous antibiotics (Megraud et al., 2021; Nestegard et al., 2022; Savoldi et al., 2018; Shu et al., 2022), in the phenotypic/functional context, we found numerous corresponding resistance genes. For instance, *cmeA, evgS, mexJ, mtrD*, and *oleC* for clarithromycin, *adeS* for tetracycline, and so on (Fig. 3C and Table S10). Presence of resistant variants of genes, for example, for erythromycin (*ermB*), quinolone (*gyrA* and *gyrB*) and tetracycline (*tetA* and *tetB*) are also known in *Campylobacter* species (Igwaran and Okoh, 2019). Bacteria may also contain inherent genes such as *AbaF*, a well-known efflux pump, that might impart antibiotic resistance, but remain functionally silent until sufficiently challenged with selection pressure (Abdi et al., 2020; Nikaido, 2009). Perturbations under antibiotics such as mutations leading to increased expression of efflux pump might result in antibiotic resistance (Nikaido, 2009; Salini et al., 2022). Antibiotic resistance can also be acquired by spontaneous mutations or through horizontal gene transfer (HGT) via plasmids (van Hoek et al., 2011). The ‘silent reservoir’ of antibiotic resistance genes can lead to the emergence of multidrug-resistant “superbugs” through HGT (Kent et al., 2020). Numerous arcobacterial members, close relatives of *H. pylori*, contain such a silent reservoir of hundreds of antibiotic resistance genes. While HGT from *H. pylori* to *C. jejuni* is known (Oyarzabal et al., 2007), *H. pylori* has a composite system for DNA uptake and natural transformation ability to possibly acquire additional antibiotic resistance (Stingl et al., 2010).

To list some of the strengths and limitations of this study – we ignored more numerous partial/incomplete genome sequences to avoid getting incomplete patterns, but this obviously reduced the number of samples for analyses. Being external tools, CARD and VFDB had their own limitations in the computational identification of target genes (Alcock et al., 2019; Liu et al., 2019) and that might have indirectly affected our analyses and results. For example, it should be noted that all the genes mentioned in this paper are listed as either virulence or antibiotic resistance genes in the respective databases based on the evidence from existing scientific literature. Finally, being purely a bioinformatics work, like others (Rao et al., 2023; Her et al., 2021), no attempts were made at experimental validations or functional characterization of these genes. However, we attempted to put the results in the phenotypic/functional context.

In conclusion, we showed that *H. pylori* genome, on average, contained 130 virulence genes – highest among *Helicobacter spp*. and 71 antibiotic resistance genes – one of the lowest among campylobacteria. While *Campylobacter spp*. showed a large difference in the number of virulence genes with *C. jejuni* containing the highest of 155, arcobacterial members showed a large difference in the number of antibiotic resistance genes and there was a positive relation with the genome size. The cags were exclusive to *H. pylori* and isolates from Asia and South America were mostly *cag2*-negative with many antibiotic resistance genes having isolate-specific patterns of occurrence. The large repository of antibiotic resistance genes along with the unique set of virulence genes might change the course of virulence and antibiotic resistance of *H. pylori* under selection pressure and in turn might pose a challenge to global public health.

## Funding and Acknowledgments

This work did not receive any specific funding. NA gratefully acknowledges the initial funding support from the OU VPRP office for the establishment of the Proteomics Core Facility.

## Statement of Ethics

The work is in compliance with ethical standards. No ethical clearance was necessary.

## Data Availability

The sequence data used in this work were obtained from NCBI. The relevant derived data are given in the supplemental result tables accessible at https://osf.io/rmd9y/.

## Author Contributions

RSPR and SDG planned and performed the work, and wrote the manuscript. LP helped in data curation. All authors contributed intellectually, and edited/reviewed the manuscript. All authors have read and agreed to the published version of the manuscript.

## Supplemental Information

Supplemental information for this article is available online.

## References

Abdi SN, Ghotaslou R, Ganbarov K, et al. (2020). Acinetobacter baumannii efflux pumps and antibiotic resistance. Infection and Drug Resistance 13:423–434.

Agresti A (2018). An introduction to categorical data analysis, 3nd ed. New York: John Wiley & Sons. 400 pages.

Alcock BP, Raphenya AR, Lau TTY, et al. (2020). CARD 2020: Antibiotic resistome surveillance with the comprehensive antibiotic resistance database. Nucleic Acids Research 48:D517–D525.

Bortolaia V, Kaas RS, Ruppe E, et al. (2020). ResFinder 4.0 for predictions of phenotypes from genotypes. Journal of Antimicrobial Chemotherapy 75:3491–3500.

Brennan DE, Dowd C, O’Morain C, et al. (2018). Can bacterial virulence factors predict antibiotic resistant Helicobacter pylori infection? World Journal of Gastroenterology 24:971–981.

Censini S, Lange C, Xiang Z, et al. (1996). Cag, a pathogenicity island of Helicobacter pylori, encodes type I-specific and disease-associated virulence□factors. Proceedings of the National Academy of Sciences USA 93:14648–14653.

Chiang T-H, Chang W-J, Chen SL-S, et al. (2021). Mass eradication of Helicobacter pylori to reduce gastric cancer incidence and mortality: A long-term cohort study on Matsu Islands. Gut 70:243–250.

Choi IJ, Kim CG, Lee JY, et al. (2020). Family history of gastric cancer and Helicobacter pylori treatment. New England Journal of Medicine 382:427–436.

Collado L, Figueras MJ (2011). Taxonomy, epidemiology, and clinical relevance of the genus Arcobacter. Clinical and Microbiological Reviews 24:174–192.

da Silva Benigno TG, Ribeiro Junior HL, de Azevedo OGR, et al. (2022). Clarithromycin-resistant H. pylori primary strains and virulence genotypes in the Northeastern region of Brazil. Revista do Instituto de Medicina Tropical de São Paulo 64:e47.

Doster E, Lakin SM, Dean CJ, et al. (2020). MEGARes 2.0: A database for classification of antimicrobial drug, biocide and metal resistance determinants in metagenomic sequence data. Nucleic Acids Research 48:D561–D569.

EFSA (2022). The European Union Summary Report on Antimicrobial Resistance in zoonotic and indicator bacteria from humans, animals and food in 2019–2020. EFSA Journal 20:7209.

Etemadi A, Safiri S, Sepanlou SG, et al. (2020). The global, regional, and national burden of stomach cancer in 195 countries, 1990–2017: A systematic analysis for the Global Burden of Disease study 2017. Lancet Gastroenterology and Hepatology 5:P4–P54.

Feldgarden M, Brover V, Gonzalez-Escalona N, et al. (2021). AMRFinderPlus and the reference gene catalog facilitate examination of the genomic links among antimicrobial resistance, stress response, and virulence. Scientific Reports 11:12728.

Ferreira EO, Lagacé-Wiens P, Klein J (2022). Campylobacter concisus gastritis masquerading as Helicobacter pylori on gastric biopsy. Helicobacter 27:e12864.

Geisinger E, Isberg RR (2017). Interplay between antibiotic resistance and virulence during disease promoted by multidrug-resistant bacteria. Journal of Infectious Diseases 215:S9–S17.

Gingold-Belfer R, Niv Y, Schmilovitz-Weiss H, et al. (2021). Susceptibility-guided versus empirical treatment for Helicobacter pylori infection: A systematic review and meta-analysis. Journal of Gastroenterology and Hepatology 36:2649–2658.

Hatakeyama M (2017). Structure and function of Helicobacter pylori CagA, the first-identified bacterial protein involved in human cancer. Proceedings of the Japan Academy B Physical and Biological Sciences 93:196–219.

Hendriksen RS, Bortolaia V, Tate H, et al. (2019) Using genomics to track global antimicrobial resistance. Frontiers in Public Health 7:242.

Her H-L, Lin P-T, Wu Y-W (2021). PangenomeNet: A pan-genome-based network reveals functional modules on antimicrobial resistome for Escherichia coli strains. BMC Bioinformatics 22:548.

Hlashwayo DF, Sigaúque B, Noormahomed EV, et al. (2021). A systematic review and meta-analysis reveal that Campylobacter spp. and antibiotic resistance are widespread in humans in sub-Saharan Africa. PLoS ONE 16:e0245951.

Hooi JKY, Lai WY, Ng WK et al. (2017). Global prevalence of Helicobacter pylori infection: Systematic review and meta-analysis. Gastroenterology 153:P420–P429.

Hosseini RS, Rahimian G, Shafigh MH, et al. (2021). Correlation between clarithromycin resistance, virulence factors and clinical characteristics of the disease in Helicobacter pylori infected patients in Shahrekord, Southwest Iran. AMB Express 11:147.

Hu Y, Zhu Y, Lu N-H (2017). Novel and effective therapeutic regimens for Helicobacter pylori in an era of increasing antibiotic resistance. Frontiers in Cellular and Infection Microbiology 7:168.

Igwaran A, Okoh, AI (2019). Human campylobacteriosis: A public health concern of global importance. Heliyon 5:e02814.

Kaakoush NO, Castaño-Rodríguez N, Mitchell HM, Man SM (2015). Global epidemiology of Campylobacter infection. Clinical Microbiology Reviews 28:687–720.

Karlsson R, Thorell K, Hosseini S, et al. (2016). Comparative analysis of two Helicobacter pylori strains using genomics and mass spectrometry-based proteomics. Frontiers in Microbiology 7:1757.

Kent AG, Vill AC, Shi Q, et al. (2020). Widespread transfer of mobile antibiotic resistance genes within individual gut microbiomes revealed through bacterial Hi-C. Nature Communications 11:4379.

Kumar S, Schmitt C, Gorgette O, et al. (2022). Bacterial membrane vesicles as a novel strategy for extrusion of antimicrobial bismuth drug in Helicobacter pylori. Mbio 13:e01633–22.

Larsson DGJ, Flach C-F (2022). Antibiotic resistance in the environment. Nature Reviews Microbiology 20:257–269.

Liu B, Pop M (2009). ARDB – Antibiotic resistance genes database. Nucleic Acids Research 37:D443–D447.

Liu B, Zheng D, Jin Q, et al. (2019). VFDB 2019: A comparative pathogenomic platform with an interactive web interface. Nucleic Acids Research 47:D687–D692.

Liu Y, Wang S, Yang F, et al. (2022). Antimicrobial resistance patterns and genetic elements associated with the antibiotic resistance of Helicobacter pylori strains from Shanghai. Gut Pathogens 14:14.

Loh JT, Shum MV, Jossart SD, et al. (2021). Delineation of the pH-responsive regulon controlled by the Helicobacter pylori ArsRS two-component system. Infection and Immunity 89:e00597–20.

Malfertheiner P, Megraud F, O’Morain CA, et al. (2017). Management of Helicobacter pylori infection – the Maastricht V/Florence Consensus Report. Gut 66:6–30.

Megraud F, Bruyndonckx R, Coenen S, et al. (2021). Helicobacter pylori resistance to antibiotics in Europe in 2018 and its relationship to antibiotic consumption in the community. Gut 70:1815–1822.

Mootapally C, Mahajan MS, Nathani NM (2021). Sediment plasmidome of the Gulfs of Kathiawar Peninsula and Arabian Sea: Insights gained from metagenomics data. Microbial Ecology 81:540–548.

Müller SA, Findeiß S, Pernitzsch SR, et al. (2013). Identification of new protein coding sequences and signal peptidase cleavage sites of Helicobacter pylori strain 26695 by proteogenomics. Journal of Proteomics 86:27–42.

Müller SA, Pernitzsch SR, Haange SB, et al. (2015). Stable isotope labeling by amino acids in cell culture based proteomics reveals differences in protein abundances between spiral and coccoid forms of the gastric pathogen Helicobacter pylori. Journal of Proteomics 126:34–45.

Nestegard O, Moayeri B, Halvorsen F-A, et al. (2022). Helicobacter pylori resistance to antibiotics before and after treatment: Incidence of eradication failure. PLoS ONE 17:e0265322.

Nikaido H (2009). Multidrug resistance in bacteria. Annual Reviews in Biochemistry 78:119–146.

Noto JM, Peek Jr RM (2012). The Helicobacter pylori cag pathogenicity island. Methods in Molecular Biology 921:41–50.

Oyarzabal OA, Rad R, Backert S (2007). Conjugative transfer of chromosomally encoded antibiotic resistance from Helicobacter pylori to Campylobacter jejuni. Journal of Clinical Microbiology 45:402–408.

Pasceri V, Cammarota G, Patti G, et al. (1998). Association of virulent Helicobacter pylori strains with ischemic heart disease. Circulation 97:1675–1679.

Pérez-Cataluña A, Salas-Massó N, Diéguez AL, et al. (2018). Revisiting the taxonomy of the genus Arcobacter: Getting order from the chaos. Frontiers in Microbiology 9:2077.

Rao RSP, Ghate SD, Shastry RP, et al. (2023). Prevalence and heterogeneity of antibiotic resistance genes in Orientia tsutsugamushi and other rickettsial genomes. Microbial Pathogenesis 174:105953.

Salini S, Muralikrishnan B, Bhat SG, et al. (2022). Overexpression of a membrane transport system MSMEG_1381 and MSMEG_1382 confers multidrug resistance in Mycobacterium smegmatis. Preprints 202204.0003.v2.

Savoldi A, Carrara E, Graham DY, et al. (2018). Prevalence of antibiotic resistance in Helicobacter pylori: A systematic review and meta-analysis in World Health Organization regions. Gastroenterology 155:1372-1382.e17.

Shu X, Ye D, Hu C, et al. (2022). Alarming antibiotics resistance of Helicobacter pylori from children in Southeast China over 6 years. Scientific Reports 12:17754.

Stingl K, Müller S, Scheidgen-Kleyboldt G, et al. (2010). Composite system mediates two-step DNA uptake into Helicobacter pylori. Proceedings of the National Academy of Sciences USA 107:1184–1189.

Suerbaum S, Michetti P (2002). Helicobacter pylori infection. New England Journal of Medicine 347:1175–1186.

Sugiyama N, Miyake S, Lin MH, et al. (2019). Comparative proteomics of Helicobacter pylori strains reveals geographical features rather than genomic variations. Genes to Cells 24:139–150.

Tacconelli E, Carrara E, Savoldi A, et al. (2018). Discovery, research, and development of new antibiotics: The WHO priority list of antibiotic-resistant bacteria and tuberculosis. Lancet Infectious Diseases 18:318–327.

Tamura K, Stecher G, Kumar S (2021). MEGA 11: Molecular evolutionary genetics analysis version 11. Molecular Biology and Evolution 10.1093/molbev/msab120.

van Hoek AHAM, Mevius D, Guerra B, et al. (2011). Acquired antibiotic resistance genes: An overview. Frontiers in Microbiology 2:203.

Vijaymeena MK, Kavitha K (2016). A survey on similarity measures in text mining. Machine Learning and Applications 3:19–28.

Wang D, Guo Q, Yuan Y, Gong Y (2019). The antibiotic resistance of Helicobacter pylori to five antibiotics and influencing factors in an area of China with a high risk of gastric cancer. BMC Microbiology 19:152.

Wei S, Li S, Wang J, et al. (2022). Outer membrane vesicles secreted by Helicobacter pylori transmitting gastric pathogenic virulence factors. ACS Omega 7:240–258.

Xu M, Liu Y, Xu C, et al. (2022). Preliminary study on the function of Cag Pathogenicity Island hp0521 gene in Helicobacter pylori 26695. https://ssrn.com/abstract=4132971.

Yuan X-y, Yan J-J, Yang Y-c, et al. (2017). Helicobacter pylori with East Asian-type cagPAI genes is more virulent than strains with Western-type in some cagPAI genes. Brazilian Journal of Microbiology 48:218–224.

Zhang Q, Zhang Z, Lu T, et al. (2020). Cyanobacterial blooms contribute to the diversity of antibiotic-resistance genes in aquatic ecosystems. Communications Biology 3:737.

Zhang Z, Zhang Q, Wang T, et al. (2022). Assessment of global health risk of antibiotic resistance genes. Nature Communications 13:1553.

