## Supplemental for "Gut feeling: Extent of virulence and antibiotic resistance genes in *Helicobacter pylori* and campylobacteria"

Running title: Virulence and antibiotic resistance genes in *H. pylori*

*Keywords: Antibiotic resistance, Chronic gastritis, Computational genomics and proteomics, Sequence analysis, Virulence*

**Note:**

The supplemental result tables (S1 to S9 and S11) are large excel files. They have been made available online and accessible at https://osf.io/rmd9y/.

**Table S1.** Basic information about *H. pylori* and campylobacterial genomes (Table_S1.xlsx).

**Table S2 and S3.** Lists of antibiotic resistance ontology (ARO, antibiotic resistance genes/mutations) in campylobacterial and *H. pylori* genomes (Table_S2.xlsx and Table_S3.xlsx – given separately due to large file sizes).

**Table S4 and S5.** Lists of virulence genes in campylobacterial and *H. pylori* genomes (Table_S4.xlsx and Table_S5.xlsx).

**Table S6 and S7.** Frequencies of antibiotic resistance ontology (ARO, antibiotic resistance genes/mutations) in campylobacterial and *H. pylori* genomes (Table_S6.xlsx and Table_S7.xlsx).

**Table S8 and S9.** Frequencies of virulence genes in campylobacterial and *H. pylori* genomes (Table_S8.xlsx and Table_S9.xlsx).

**Table S10.** List of known antibiotic resistance in *H. pylori* and likely resistance genes.

| Antibiotic | Class | MIC (μg/ml) | Genes | Reference |
| --- | --- | --- | --- | --- |
| Amoxicillin | Amoxicillin | ≥ 0.5 | *H. pylori pbp1, H. pylori pbp2, H. pylori pbp3* | Nishizawa et al., 2011 |
| Chloramphenicol | Chloramphenicol | ≥ 4 | *acrB, catB10, catI, cfr(D), cipA, clcD, cmlv, mexN, mexV, optrA, parS, poxtA, R. fascians cmr, rsmA, YajC* | Kwon et al., 2003 |
| Ciprofloxacin | Fluoroquinolones | ≥ 8 | *acrB, arlR, C. difficile gyrA, C. difficile gyrB, cdeA, H. pylori gyrA, H. pylori gyrB, hp1181, mdtK, mexI, patA, pmpM, rsmA, yajC* | Garcia et al., 2012 |
| Clarithromycin | Macrolides | ≥ 1 | *carA, cmeA, efrB, erm(30), erm(34), erm(44)v, erm(50), erm(51), ermB, ermG, ermY, evgS, lpeB, macB, mexJ, mexV, mreA, mtrA, mtrD, oleC, parS, srmB, tlrC* | Nishizawa et al., 2011 |
| Clindamycin | Lincosamide | ≥ 256-512 | *cfr(D), cipA, clcD, erm(30), erm(34), erm(44)v, erm(50), erm(51), ermB, ermG, ermY, ImrD, IsaC, lmrC, salA, salC, salD, salE, tlrC, vgaA, vgaALC, vmlR* | Wang and Taylor, 1998 |
| Levofloxacin | Fluoroquinolones | ≥ 32 | *acrB, arlR, C. difficile gyrA, C. difficile gyrB, cdeA, H. pylori gyrA, H. pylori gyrB, hp1181, mdtK, mexI, patA, pmpM, rsmA, yajC* | Li et al., 2021 |
| Metronidazole | Nitroimidazole | ≥ 8 | *H. pylori frxA, H. pylori rdxA* | Mehrotra et al., 2021 |
| Quinupristin | Streptogramins | ≥ 64-128 | *cfr(D), cipA, clcD, erm(30), erm(34), erm(44)v, erm(50), erm(51), ermB, ermG, ermY, lsaC, salA, salC, salD, salE, tlrC, vatA, vatC, vatF, vatH, vgaA, vgaALC, vgaB, vmlR* | Wang and Taylor, 1998 |
| Rifampicin | Rifampicin | > 1 | *B.adolescentis rpoB, C. difficile rpoB, H. pylori rpoB, M. tuberculosis rpoB, rphB, S. aureus rpoB* | Tang et al., 2022 |
| Tetracycline | Tetracycline | ≥ 4 | *adeB, adeR, adeS, mepA, poxtA, S. rimosus otr(A), tet(35), tet(36), tet(44), tet(M), tet(S), tet(T), tet(W/32/O), tetA(46), tetA(58), tetA(60), tetA(P), tetB(46), tetB(60), TxR* | Ribeiro et al., 2004 |
| Vancomycin | Glycopeptide | > 10 | *vanA_H, vanB_R, vanC, vanE, vanE_R, vanF_H, vanF_R, vanG_T, vanL, vanL_S, vanL_Tr, vanM_H, vanN_R, vanP_S* | Kusters et al., 2006 |

**References**

Garcia M, Raymond J, Garnier M, et al. (2012). Distribution of spontaneous gyrA mutations in 97 fluoroquinolone-resistant *Helicobacter pylori* isolates collected in France. Antimicrobial Agents and Chemotherapy 56:550-551.

Kusters JG, Van Vliet AH, Kuipers EJ (2006). Pathogenesis of *Helicobacter pylori* infection. Clinical microbiology reviews 19:449-490.

Kwon D H, Dore MP, Kim JJ, et al. (2003). High-level β-lactam resistance associated with acquired multidrug resistance in *Helicobacter pylori*. Antimicrobial agents and chemotherapy 47:2169-2178.

Li J, Deng J, Wang Z, et al. (2021). Antibiotic resistance of *Helicobacter pylori* strains isolated from pediatric patients in southwest China. Frontiers in Microbiology 11:621791.

Mehrotra T, Devi TB, Kumar S, et al. (2021). Antimicrobial resistance and virulence in *Helicobacter pylori*: Genomic insights. Genomics 113:3951-3966.

Nishizawa T, Suzuki H, Tsugawa H, et al. (2011). Enhancement of amoxicillin resistance after unsuccessful *Helicobacter pylori* eradication. Antimicrobial agents and chemotherapy 55:3012-3014.

Ribeiro ML, Gerrits MM, Benvengo YH, et al. (2004). Detection of high-level tetracycline resistance in clinical isolates of *Helicobacter pylori* using PCR-RFLP. FEMS Immunology & Medical Microbiology 40:57-61.

Tang X, Wang Z, Shen Y, et al. (2022). Antibiotic resistance patterns of *Helicobacter pylori* strains isolated from the Tibet Autonomous Region, China. BMC Microbiology 22:196.

Wang GE, Taylor DE (1998). Site-specific mutations in the 23S rRNA gene of *Helicobacter pylori* confer two types of resistance to macrolide-lincosamide-streptogramin B antibiotics. Antimicrobial Agents and Chemotherapy 42:1952-1958.

**Table S11.** Lists of *H. pylori* proteins identified by various proteomic studies (Table_S11.xlsx).

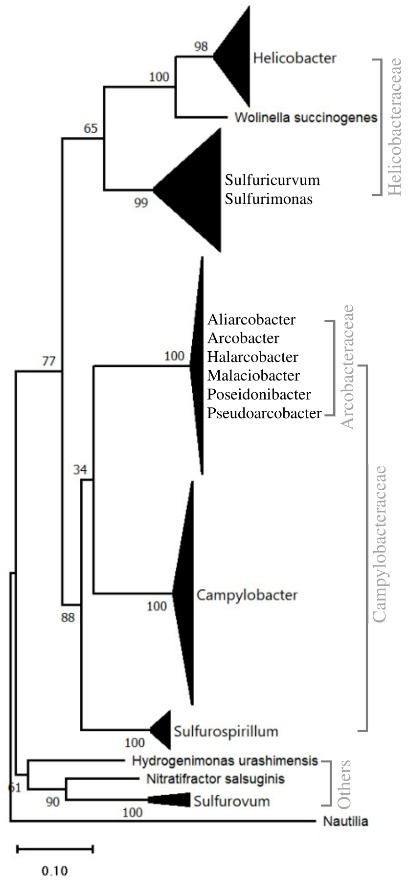

**Fig. S1.** Phylogeny of campylobacteria. The tree was based on 120 non-redundant (out of 316) complete 16S rRNA sequences from 91 species of campylobacteria. *Nautilia profundicola* was taken as the outgroup. Numbers next to nodes indicate bootstrap values (%) based on 1000 iterations. Branch length scale indicates the number of substitutions per site. The phylogeny tree was constructed in MEGA11 using the maximum likelihood method.

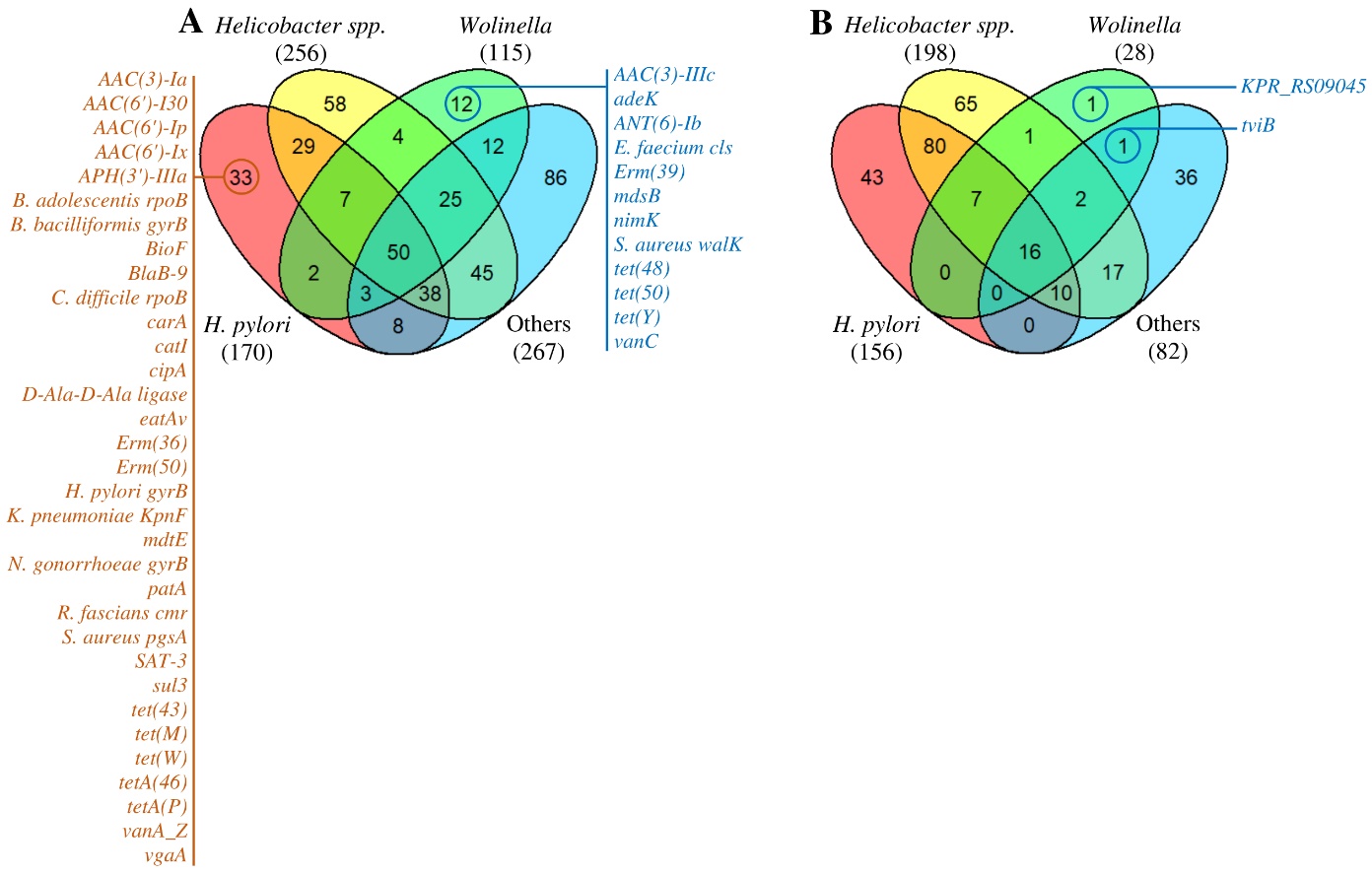

**Fig. S2.** Venn diagrams showing the overlap in the presence of (A) antibiotic resistance genes and (B) virulence genes in the Helicobacteraceae clades. (A) Numerous antibiotic resistance genes are unique to each clade within the Helicobacteraceae, for example, 12 genes in *Wolinella*, a close relative of *Helicobacter*, and 86 genes in others (*Sulfuricurvum* and *Sulfurimonas spp.*). (B) *Wolinella* and others (*Sulfuricurvum* and *Sulfurimonas spp.*) have far fewer virulence genes with just one unique gene in the former and 36 in the latter. Note: *H. pylori* was not included in *Helicobacter spp.* set.

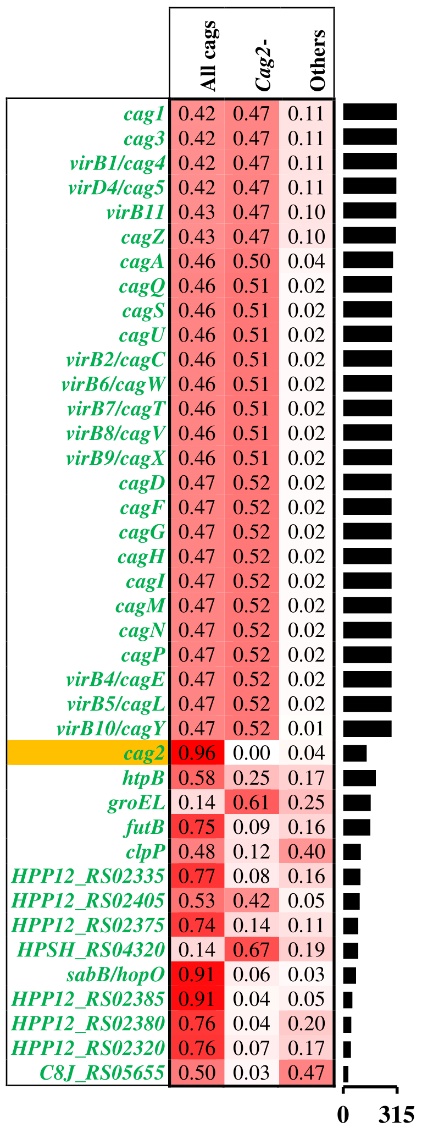

**Fig. S3.** Heat map shows the relative proportion of genes in three groups – apart from all cags, a few more virulence genes such as *htpB*, *groEL*, etc. were also significantly different (p < 0.05, chi-squared test with Bonferroni correction) in one of the groups based on cags. Bar graph at the right shows the number of genomes.

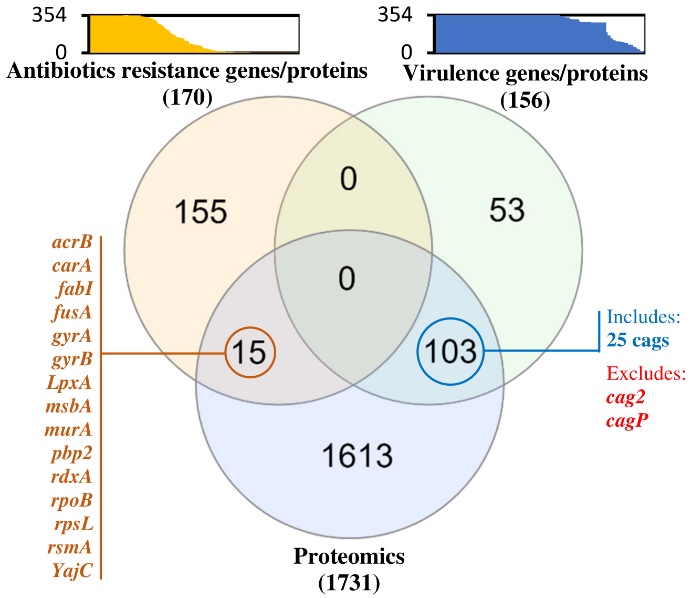

**Fig. S4.** Just 15 (8.8%) antibiotic resistance genes, but 103 (66.0%) virulence genes were proteomically identified based on *H. pylori* proteomic studies. Bar charts at top panel show the frequency of genes – only 67 (39.4%) antibiotic resistance genes but 128 (82.1%) virulence genes were present in 50% or more of genomes.
